# Antigenic and Virological Characteristics of SARS-CoV-2 Variant BA.3.2, XFG, and NB.1.8.1

**DOI:** 10.1101/2025.04.30.651462

**Authors:** Caiwan Guo, Yuanling Yu, Jingyi Liu, Fanchong Jian, Sijie Yang, Weiliang Song, Lingling Yu, Fei Shao, Yunlong Cao

## Abstract

The emergence of the SARS-CoV-2 saltation variant BA.3.2, which harbors over 50 mutations relative to its ancestral BA.3 lineage, has raised concerns about its potential to drive outbreaks similar to BA.2.86/JN.1. Concurrently, variants such as NB.1.8.1, LF.7.9, XEC.25.1, XFH, and XFG exhibit enhanced growth advantages over LP.8.1.1, necessitating a comparative analysis of their antigenic and virological characteristics. Here, we evaluated the infectivity, ACE2-binding efficiency, and immune evasion of these variants. Pseudovirus assays revealed BA.3.2’s robust antibody evasion, including resistance to Class 1/4 monoclonal antibodies; however, its ACE2 engagement efficiency was markedly reduced due to a closed spike conformation, leading to low infectivity. While XFG and LF.7.9 demonstrated strong immune escape associated with A475V and N487D mutations, their reduced receptor-binding efficiency suggested a need for compensatory adaptations. In contrast, NB.1.8.1 retained high ACE2 affinity and humoral immune evasion, supporting its potential for future dominance. Collectively, BA.3.2’s current profile limits its ability to compete with emerging variants like NB.1.8.1. Sustained monitoring of BA.3.2’s evolution— particularly for mutations stabilizing an open RBD conformation or enhancing escape from Class 1 antibodies—is essential to assess its outbreak potential.

## Main Text

The SARS-CoV-2 saltation variant BA.3.2, harboring over 50 mutations relative to its ancestral BA.3 lineage, has recently drawn global attention (Fig. 1A). Notably, BA.3.2 exhibits 44 mutations distinct from the currently dominant LP.8.1/LP.8.1.1 variant (Fig. S1), raising speculation about its potential to drive an outbreak similar to BA.2.86/JN.1, particularly following its first detection outside South Africa in the Netherlands on April 2, 2025.^1-5^ A critical evaluation of its antigenic profile and infectivity is essential to determine its likelihood of prevailing.

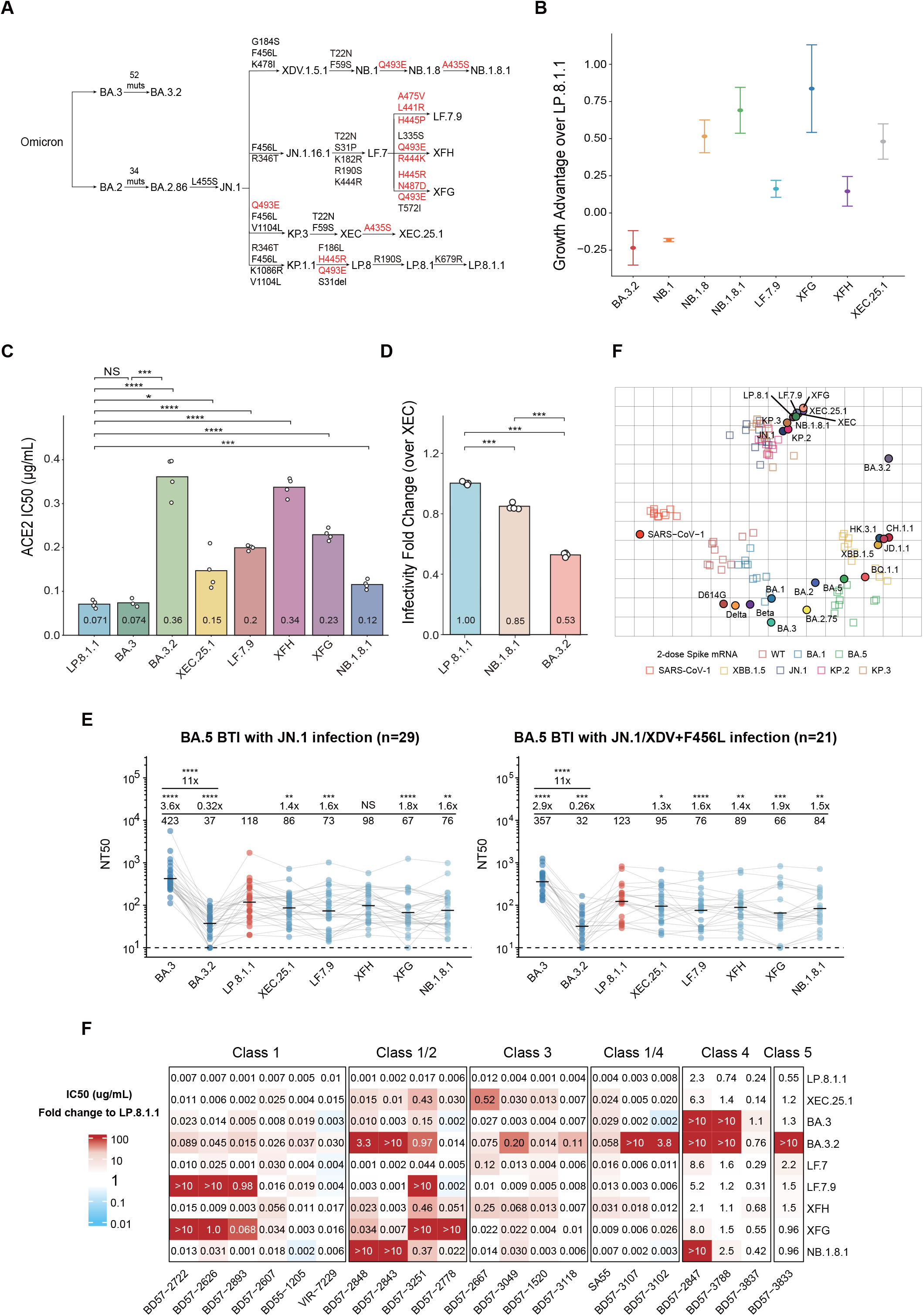
(A)Genetic changes in the spike glycoprotein of prevalent SARS-CoV-2 variants. Key mutations observed in these variants are marked in red. (B) The relative growth advantage of BA.3.2, NB.1, NB.1.8, NB.1.8.1, LF.7.9, XFG, XFH, XEC.25.1, compared with LP.8.1.1. Daily global sequence data retrieved from the GISAID database (April 1, 2024 - April 11, 2025) were used to compute the relative growth advantage. Growth advantage was calculated with the generation time set to 7 days, and confidence intervals were calculated with α = 0.95. The symbol ‘*’ signifies all sublineages. (C) IC_50_ values for the neutralization of LP.1.8.1, BA.3, BA.3.2, XEC.25.1, LF.7.9, XFG, XFH, and NB.1.8.1 pseudoviruses by soluble human ACE2. Technical replicates are shown as individual circles. IC_50_ (μg/mL) was log-transformed prior to two-tailed t test analysis. (D) Relative infectivity of NB.1.8.1 and BA.3.2 compared to XEC in Vero cells. Infectivity was assessed using vesicular stomatitis virus (VSV) pseudoviruses. Each dot represents a replicate and bars denote mean values. Statistical significance was evaluated using two-tailed t test. (E) Antigenic map based on NT_50_ titers from plasma of mice immunized with two doses of spike mRNA vaccines against indicated strains. The map was generated using modified multidimensional scaling (MDS) in Racmacs (v1.1.35) and visualized with ggplot2 (v3.4.1) after 500 optimization steps with the minimum column basis set to “none”. (F) NT_50_ values were measured from convalescent plasma of individuals reinfected with JN.1 post BA.5/BF.7 breakthrough (n=29), or with JN.1/XDV (Phe456Leu) post BA.5/BF.7 (n=21). Cohort labels and sample sizes are provided above each panel. The dashed line represents the detection limit (NT_50_ = 10). Geometric means, fold reductions, and corresponding p values (Wilcoxon rank-sum test) are noted above each group. (G) IC_50_ values (μg/mL) of monoclonal neutralizing antibodies targeting RBD epitopes against BA.3, BA.3.2, LP.8.1.1, XEC.25.1, LF.7.9, XFH, XFG, NB.1.8.1 variants. Fold changes relative to LP.8.1.1 are shown by background color; red denotes increased resistance, blue denotes enhanced sensitivity. Color scale indicates the fold-change magnitude.

Concurrently, multiple emerging variants—including NB.1.8.1, LF.7.9, XEC.25.1, XFH, and XFG—exhibit enhanced growth advantages over LP.8.1.1, suggesting their potential to dominate future transmission waves (Fig. 1B). These variants demonstrate convergent evolution of recurrent mutations such as Q493E, A435S, and A475V (Fig. 1A).^6-8^ Specifically, NB.1.8 and NB.1.8.1, descendants of the XDV.1.5.1 sublineage, are characterized by Q493E and A435S mutations, respectively, and have rapidly spread in China and Hong Kong. Similarly, XEC.25.1, a derivative of XEC, harbors the A435S mutation and also demonstrates a high growth advantage. LF.7.9, a highly fit European variant derived from LF.7, carries the receptor-binding domain (RBD) mutations A475V, L441R, and H445P. The recombinant XFG variant, originating from LF.7 and LP.8.1.2, harbors four critical spike mutations (H445R, N487D, Q493E, T572I) and has achieved rapid global spread following its initial detection in Canada. Another recombinant variant, XFH, originating from LF.7.1 and XEF, carries the convergent mutation Q493E, a reversion of R444K, and a novel L335S mutation, also positioning it among the fastest-growing variants to date. The concurrent emergence of BA.3.2 alongside these variants necessitates urgent comparative analysis of their antigenic and virological characteristics to evaluate whether BA.3.2 can outcompete existing lineages.

First, we generated spike-pseudotyped vesicular stomatitis viruses for the concerning variants and assayed the efficiency of soluble human ACE2 (hACE2) to inhibit viral entry, which reflects the ACE2-binding strength and receptor engagement efficiency of each variant’s spike protein (Fig. 1C). Recombinant RBD subunits were also produced, and their binding affinity to hACE2 was quantified via surface plasmon resonance (Fig. S2). Surprisingly, all tested variants exhibited relatively lower ACE2-binding capability compared to LP.8.1.1. Specifically, BA.3.2’s spike exhibited the lowest ACE2 engagement efficiency, significantly reduced compared to its ancestor BA.3 and LP.8.1.1. However, BA.3.2’s RBD actually displayed similar ACE2-binding affinity to LP.8.1.1, suggesting that its reduced spike-mediated ACE2 engagement arises from a relatively “closed” spike or “down” RBD conformation. Similarly, XFH exhibited low ACE2 engagement efficiency, likely due to a closed spike conformation induced by the L335S mutation. In contrast, LF.7.9 and XFG showed markedly reduced RBD-ACE2 binding affinity, attributable to their A475V and N487D mutations, respectively, which explain their lower receptor engagement efficiency. Notably, XEC.25.1 and NB.1.8.1 retained robust ACE2 engagement, with NB.1.8.1 exhibiting the highest RBD-ACE2 binding affinity among all variants tested. Consistent with these findings, pseudovirus infectivity assayed in Vero cells revealed that BA.3.2 exhibits disastrously low fitness compared to LP.8.1.1, while NB.1.8.1 retains acceptable infectivity (Fig. 1D).

Next, we evaluated the antigenicity and humoral immune evasion properties of these variants using pseudovirus neutralization assays. Antigenicity was assessed using serum samples from naïve mice immunized with two doses of spike mRNA vaccines, while antibody evasion capability was tested with human convalescent plasma. The plasma used in this study was obtained from two cohorts of individuals who received 2–3 doses of inactivated SARS-CoV-2 vaccines and subsequently experienced BA.5 or BF.7 breakthrough infections, with one cohort reinfected by JN.1 (n=29) and the other by JN.1 or XDV with F456L (n=21), as previously described.^3^ Notably, BA.3.2 exhibited high resistance to serum neutralization across all naïve mouse vaccine groups (Fig. 1D). Antigenic cartography based on pseudovirus neutralization titers revealed BA.3.2 to be antigenically distinct from the JN.1 and XBB.1.5 lineages, whereas LF.7.9, XEC.25.1, XFG, and NB.1.8.1 all clustered close to the JN.1 family (Fig. 1D).^8^ In human plasma, BA.3.2 also demonstrated profound humoral immune evasion, with an 11-fold reduction in geometric mean neutralizing titer (NT50) compared to BA.3 and a 3–4-fold reduction relative to LP.8.1.1 (Fig. 1E). While BA.3.2 showed the strongest evasion, XFG, LF.7.9, and NB.1.8.1 also displayed enhanced escape compared to LP.8.1.1 in both cohorts: XFG exhibited a nearly 2-fold decrease in NT50, whereas LF.7.9 and NB.1.8.1 showed 1.5–1.6-fold reductions, consistent with their relative growth advantages over LP.8.1.1 (Fig. 1E).

To delineate the molecular basis of immune evasion, we profiled the sensitivity of these variants to a panel of RBD-targeting neutralizing monoclonal antibodies (mAbs) spanning distinct epitopes (Fig. 1F and S3). LF.7.9 exhibited pronounced resistance to Class 1 antibodies, primarily driven by its A475V mutation, while XFG resistance was attributed to its N487D and Q493E mutations. The N487D mutation in XFG additionally conferred escape from Class 1/2 (Group B) antibodies.^8^ Similarly, the K478I mutation in NB.1.8.1 and K478N in BA.3.2 enhanced the evasion of Class 1/2 antibodies. The A435S mutations in XEC.25.1 and NB.1.8.1 reduced antibody neutralization potency across all epitopes, similar to observations in MC.10.1.^3^ Strikingly, BA.3.2 demonstrated robust escape from Class 1/4 antibodies, a class of potent, broad-spectrum neutralizing antibodies effective against most Omicron lineages, including LF.7.9, XEC.25.1, XFG, XFH, and NB.1.8.1 (Fig. 1E). Notably, Class 1/4 antibodies are prevalent in Chinese populations immunized with inactivated vaccines, where immune imprinting is less pronounced.^9,10^ In contrast, mRNA vaccine recipients with stronger immune imprinting rarely develop these antibodies, which may cause divergent neutralization responses against BA.3.2 between these groups.

In summary, our findings indicate that BA.3.2 exhibits robust antibody evasion but suffers from low ACE2-binding capability and infectivity, significantly limiting its likelihood of prevailing. To achieve efficient spread akin to BA.2.86 or JN.1, BA.3.2 would require additional mutations to improve both its receptor engagement efficiency (e.g., stabilizing an “open” RBD conformation) and its evasion of Class 1 antibodies. Similarly, while XFG displays strong immune evasion, its relatively low ACE2 engagement efficiency suggests it may need compensatory mutations to enhance receptor compatibility for sustained transmission. Importantly, NB.1.8.1 demonstrates a balanced profile of ACE2 binding and immune evasion, supporting its potential for future prevalence.

## Declaration of interests

Y.C. has provisional patent applications for the BD series antibodies (WO2024131775A9 and WO2023151312A1), and is the founder of Singlomics Biopharmaceuticals. The other authors declare no competing interests..

## Appendix

## Acknowledgments

We appreciate the persistent efforts of the scientific community in monitoring SARS-CoV-2 variants and extend our thanks to all the volunteers who provided blood samples. This study was jointly supported by National Science and Technology Major Project (2022ZD0115002), Changping Laboratory (2021A0201; 2021D0102), and the National Natural Science Foundation of China (32222030, 2023011477).

## Author Contributions

Y.C. designed and supervised the study. C.G. and Y.C. wrote the manuscript with inputs from all authors. C.G., J.L., S.Y., F.J., and W.S. performed sequence analysis, illustration, and figure preparation. Y.Y. constructed pseudoviruses. L.Y., and F.S. processed the plasma samples and performed the pseudovirus neutralisation assays. F.J. and Y.C. analyzed the neutralisation data.

## Methods Details

### Growth Advantage Calculation

An algorithm adapted from Chen et al^1^. was used to quantify growth advantages based on global SARS-CoV-2 sequences from the GISAID database^2^ collected between April 1, 2024 and April 11, 2025. A logistic regression model was fitted to the daily sample frequency of concerning strains BA.3.2, NB.1, NB.1.8, NB.1.8.1, LF.9, LF.7.2.1, LF.7.7.2, XFG, XFH and XEC.25.1 to estimate each strain’s logistic growth rate (a) and sigmoid-curve midpoint (t_0_). Growth advantage (e ^a × g^ −1) was calculated with g set to 7 days (the generation time), and confidence intervals were calculated with α = 0.95. To determine the relative growth advantage compared to LP.8.1.1, the total number of sequences for the target variant and the reference strains was used as the background.

### Surface Plasmon Resonance

Surface plasmon resonance (SPR) analysis was performed by Biacore 8K (Cytiva). Human ACE2 protein was digested for 1 hour at 37°C with 5% CO_2_, followed by immobilization on Protein A sensor chips (Cytiva, 29127556). Purified SARS-CoV-2 variant RBD protein (His-tagged, Sino Biological) was diluted in a series of gradient concentrations (6.25, 12.5, 25, 50, and 100 nM) and injected onto the chips. Binding data were collected at room temperature using Biacore 8K Control Software (v4.0.8.20368, Cytiva), and kinetic parameters were analyzed with the 1:1 binding model using Biacore 8K Evaluation Software (v4.0.8.20368, Cytiva). Each variant was performed with at least two independent replicates to ensure data reliability.

### Patient recruitment and plasma isolation

Volunteers who experienced reinfection with the SARS-CoV-2 Omicron BTI variant provided blood samples. The research protocol received the approval from the Ethics Committees of Beijing Ditan Hospital Capital Medical University (Ethics Committee Archiving No. LL-2021-024-02), the Tianjin Municipal Health Commission, and the ethics committee of Tianjin First Central Hospital (Ethics Committee Archiving No. 2022N045KY). All participants submitted written informed consent following the Declaration of Helsinki, granting approval for the collection, storage, and research use of clinical samples including the release of related data.

Volunteers were patients initially infected in Beijing and Tianjin during December 2022, when the infections were predominantly caused by BA.5* variants. From December 2022 to February 2023, sequencing of clinical samples revealed that >98% were identified as the BA.5/BF.7 lineage, primarily composed of BA.5.2.48 and BF.7.14 subvariants. We defined two reinfection cohorts. The JN.1 reinfection cohort experienced their initial breakthrough infections after receiving two to three doses of inactivated vaccines, and reinfected between February and March 2024. The JN.1/XDV+F456L reinfection cohort consists of volunteers who contracted their initial breakthrough infections after completing a three-dose inactivated vaccine, followed by reinfecting between July and August 2024, when 97 % of sequenced samples were identified as JN.1+F456L or XDV+F456L variants. All infections were confirmed through PCR or antigen testing.

The whole blood was diluted 1:1 in PBS containing 2 % fetal bovine serum, and then peripheral blood mononuclear cells (PBMCs) and plasma were isolated through density-gradient centrifugation using Ficoll (Cytiva, 17-1440-03). After centrifugation, plasma samples in the upper layer were collected, aliquoted and stored at −20 °C or lower. All the plasma samples underwent heat inactivation before experimental use.

### Pseudovirus neutralisation assay

We generated SARS-CoV-2 variant spike pseudoviruses by using the Vesicular stomatitis virus (VSV) pseudovirus packaging system. G*ΔG-VSV (VSV-G pseudotyped virus, Kerafast) and spike-expressing plasmid were transfected to 293T cells (American Type Culture Collection [ATCC], CRL-3216). After cell culture, the pseudovirus in the supernatant was collected, filtered, aliquoted, and stored at −80 °C for future use.

For the neutralization assay, serial dilutions of monoclonal antibodies, human ACE2-Fc, and plasma samples were mixed with prepared pseudovirus in 96-well plates and the mixtures were incubated at 37°C with 5% CO_2_ for 1 hour. Plasma samples were diluted at 1:10 initially, then performed 3-fold serial dilutions for 5 times. Monoclonal anti-S-RBD antibodies and ACE2-Fc were diluted to 0.1 mg/mL at first, then diluted 50-fold, followed by five 5-fold serial dilutions. Digested Huh-7 cells (Japanese Collection of Research Bioresources [JCRB], 0403) were then seeded to each well. After 24 hours of incubation, the supernatant was removed, and D-luciferin reagent (vazyme, DD1209-03) was added. The reaction lasted for 2 minutes in the dark, after which Luminescence was measured by using a microplate spectrophotometer (PerkinElmer HH3400). The half-maximal inhibitory concentration (IC_50_) values were calculated using a four-parameter logistic regression model.

## Supplementary Figures and Tables

**Fig. S1.**
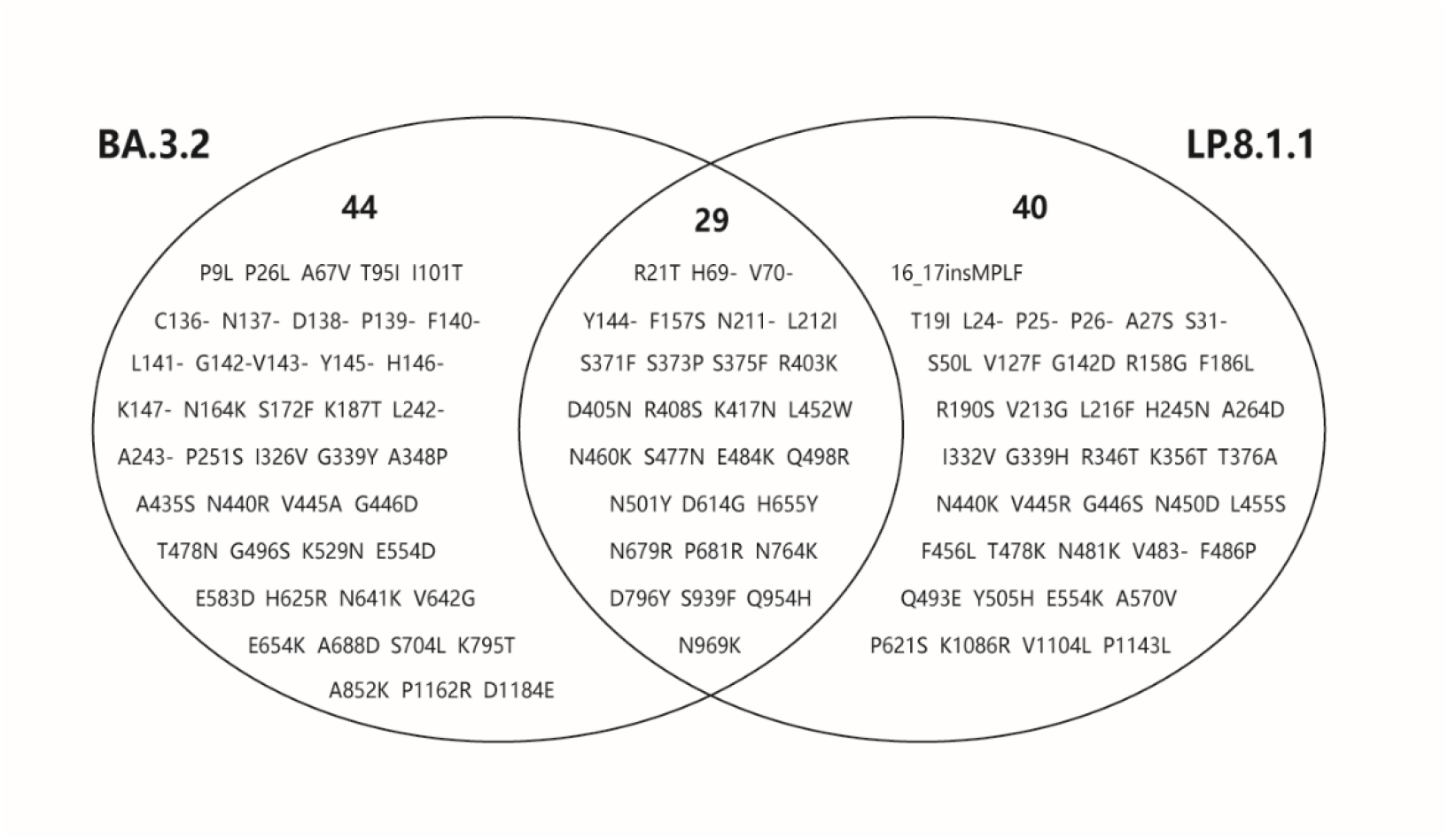
Venn Diagram of Shared and Unique Mutations between BA.3.2 and LP.8.1.1.

**Figure S2.**
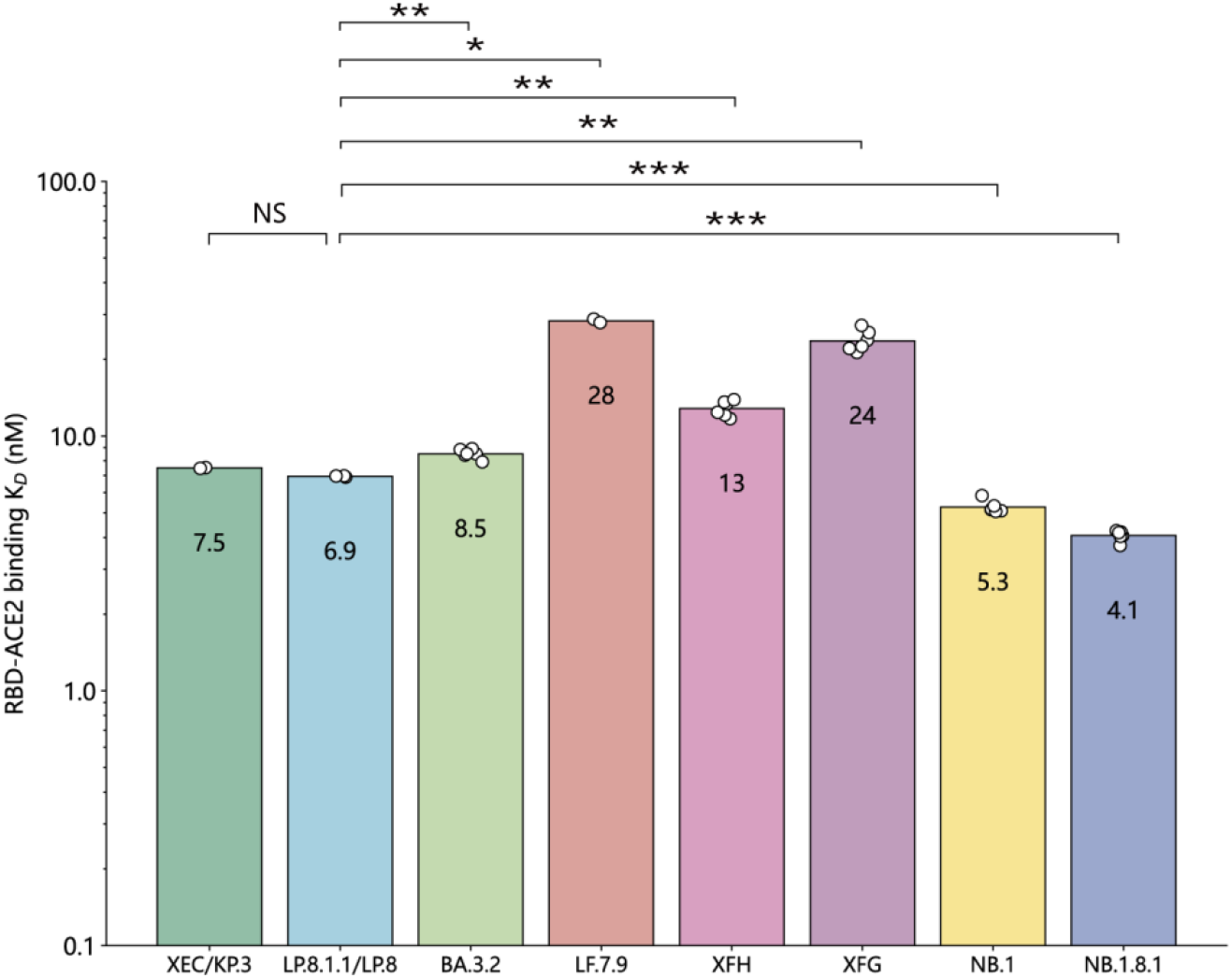
The Binding Affinity of KP.3/XEC, LP.8.1.1/LP.8, BA.3.2, LF.9, XFH, XFG, NB.1, and NB.1.8.1 RBD Proteins to Human ACE2. The binding affinity of RBD proteins to human ACE2, established by SPR. Each circle indicates a technical replicate. Geometric mean KD values (nM) are displayed. Log-transformed data were analysed using a two-tailed t-test to compare the means between the two groups. SPR=surface plasmon resonance.

**Figure S3.**
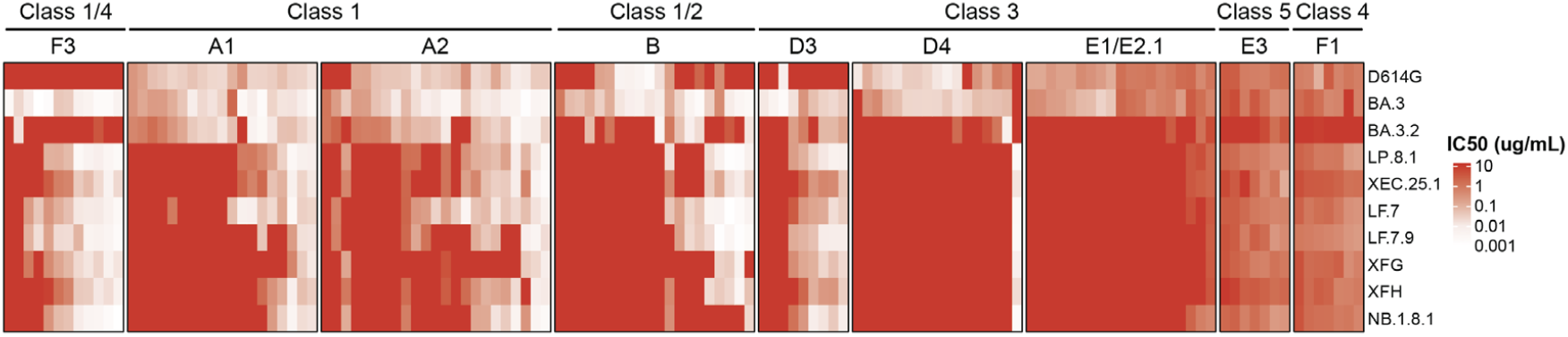
Escaping profiles of recent SARS-CoV-2 variants against an antibody panel targeting distinct RBD epitopes. IC_50_ values for a panel of monoclonal neutralising antibodies targeting RBD epitopes against SARS-CoV-2 variants. The values within the table are IC50 values (μg/mL), while the background colour indicates the fold-change in IC50 relative to LP.8.1.1. The colour gradient bar to the right represents the magnitude of IC50 fold-change.

**Table S1.**
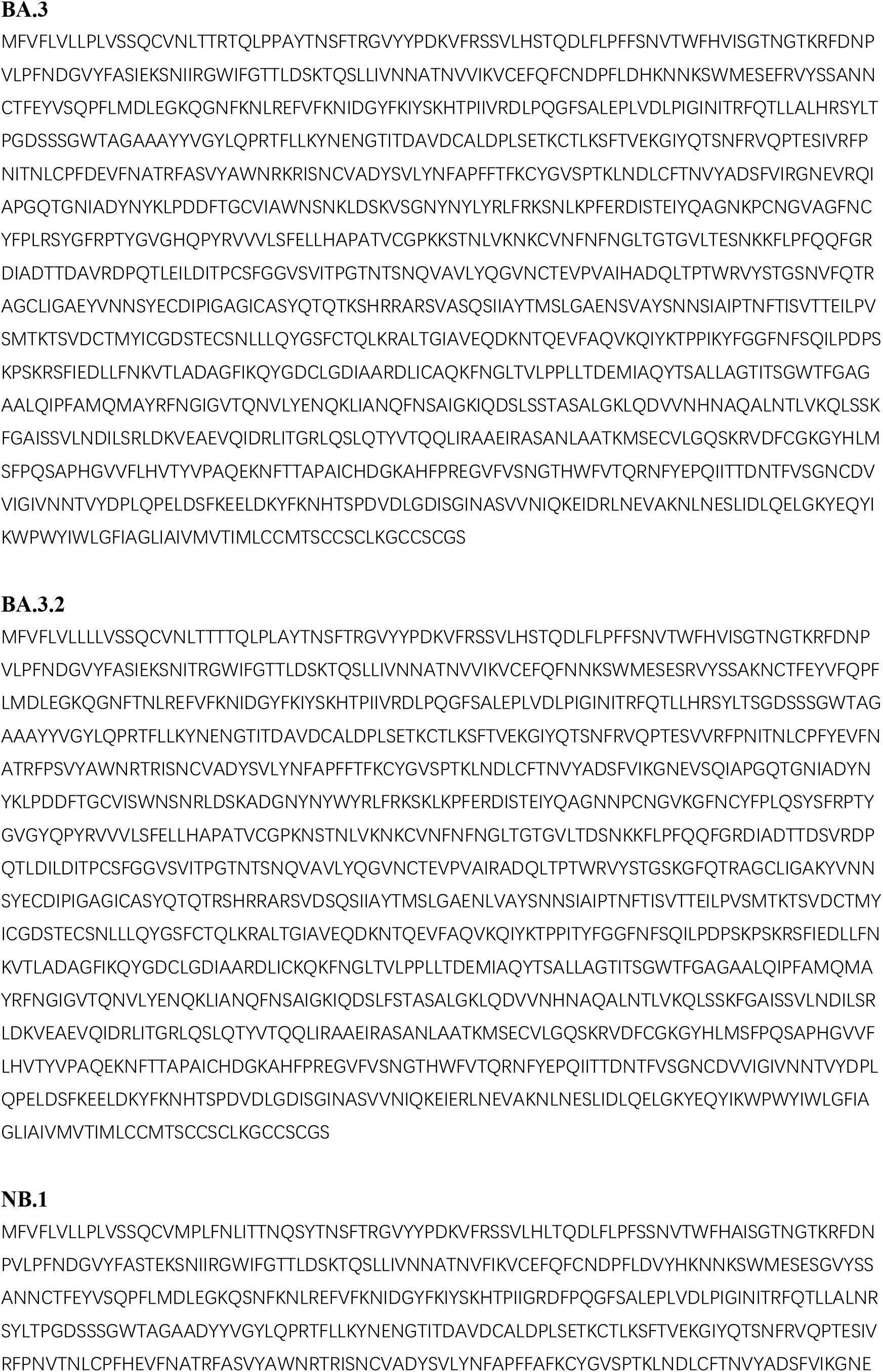
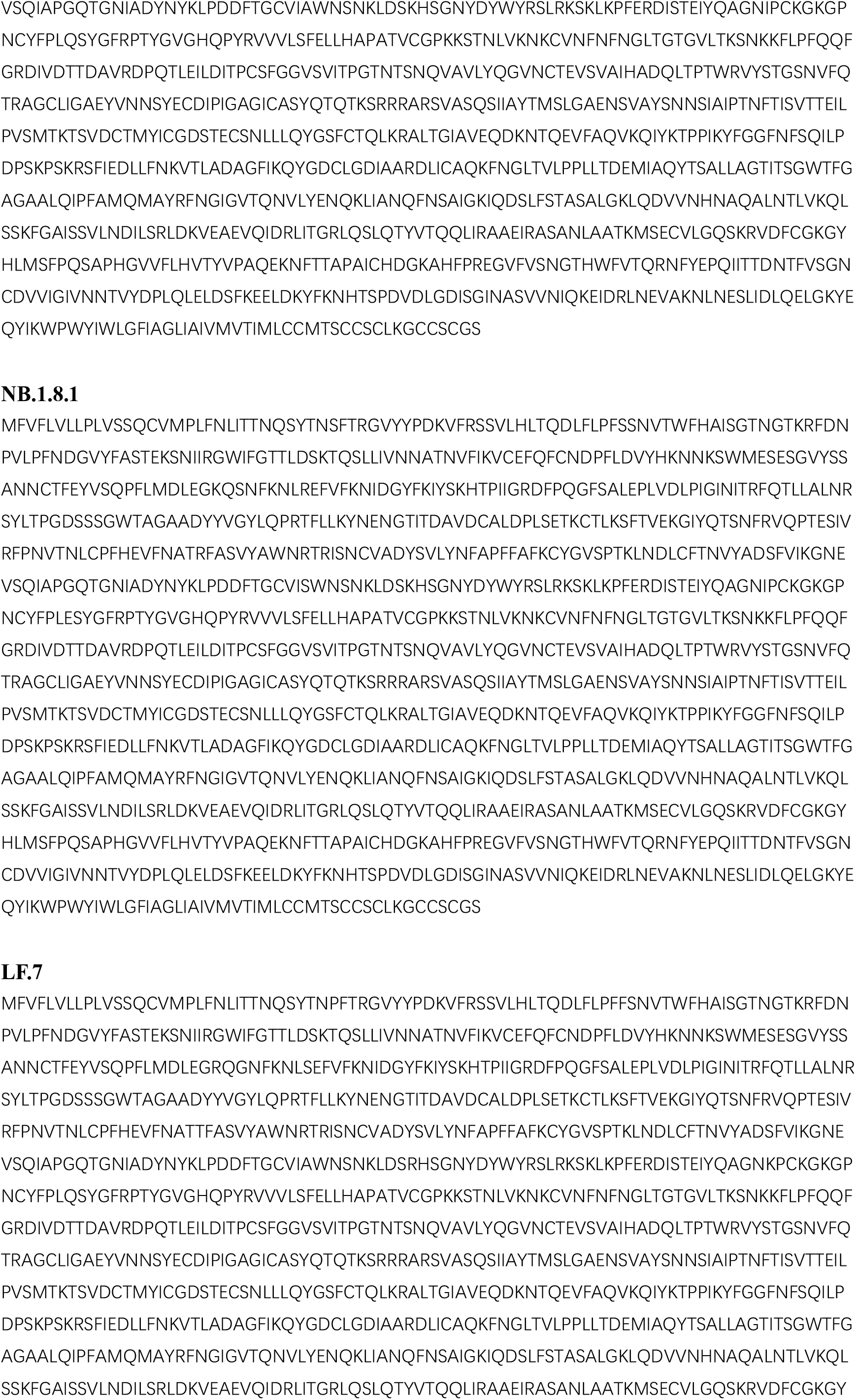
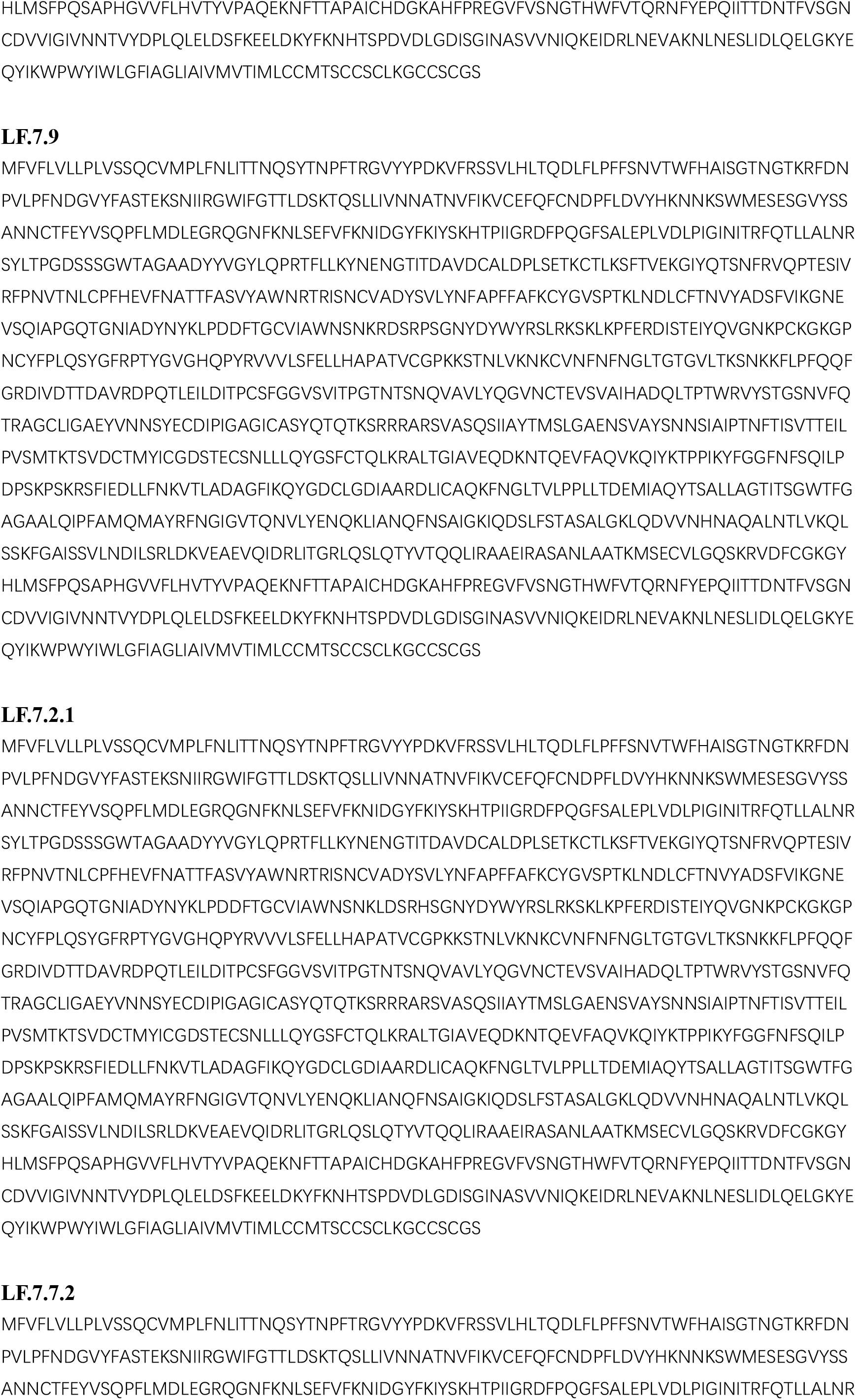
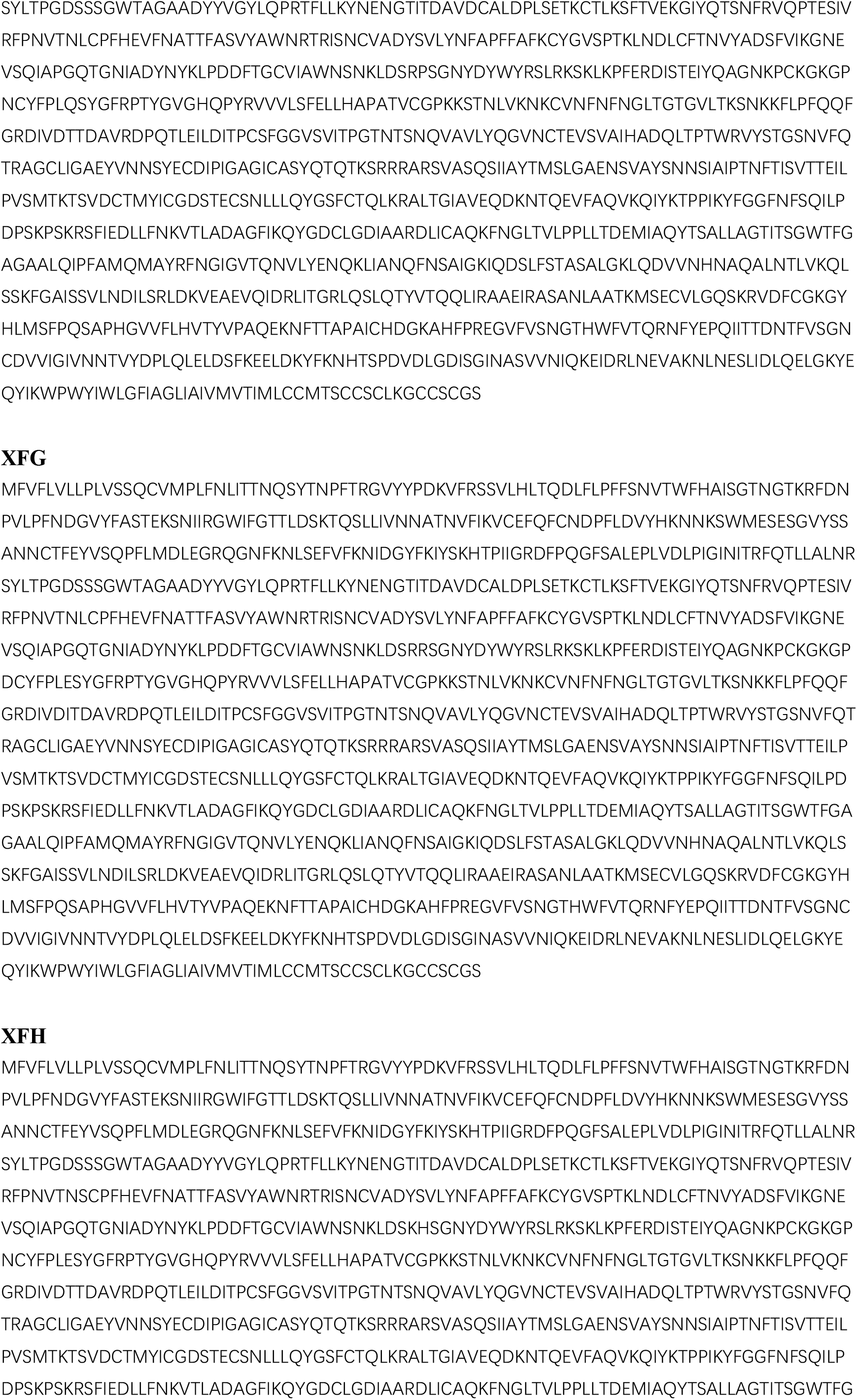
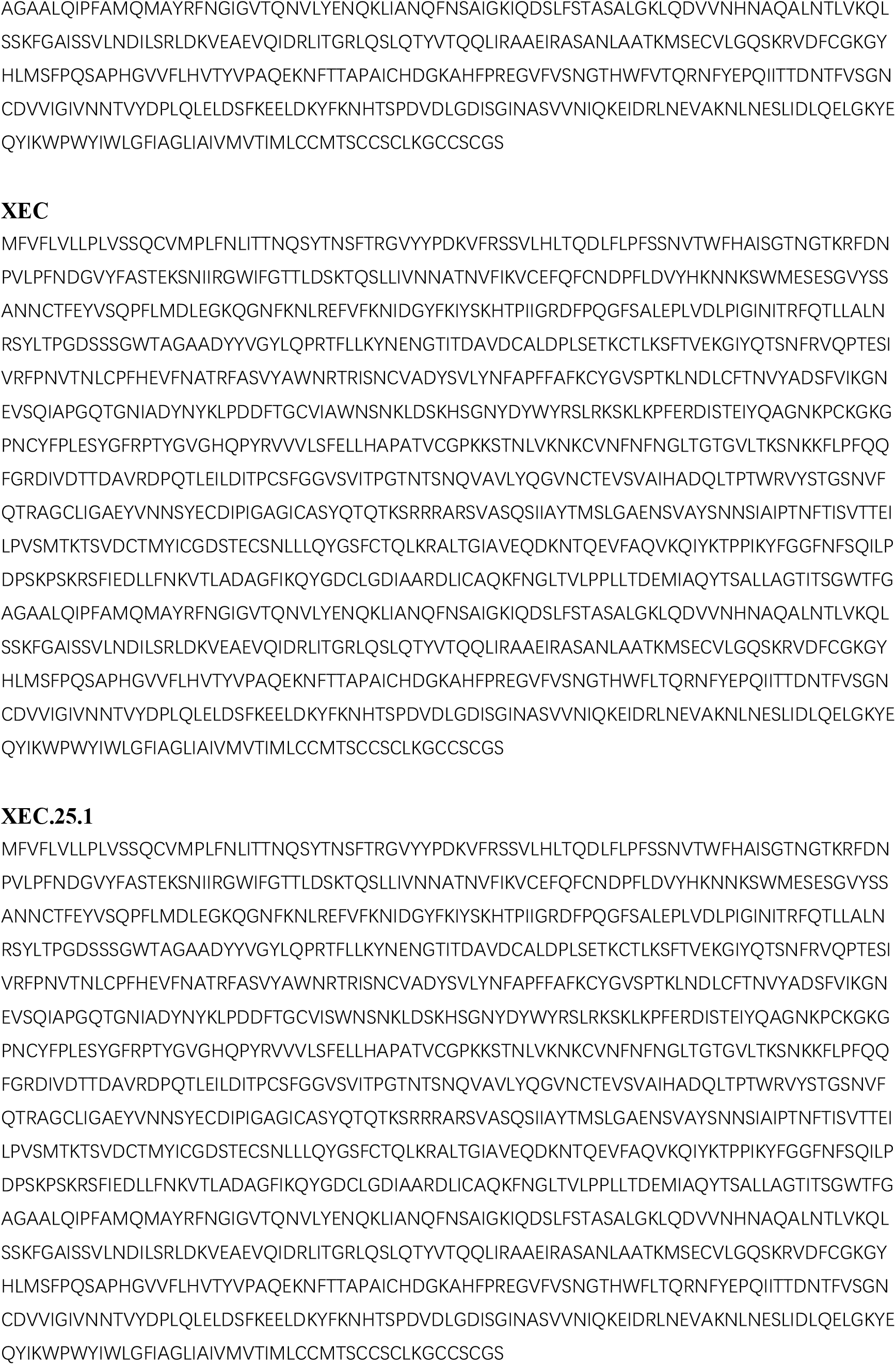
Spike protein sequences of SARS-CoV-2 variants.

**Table S2.**
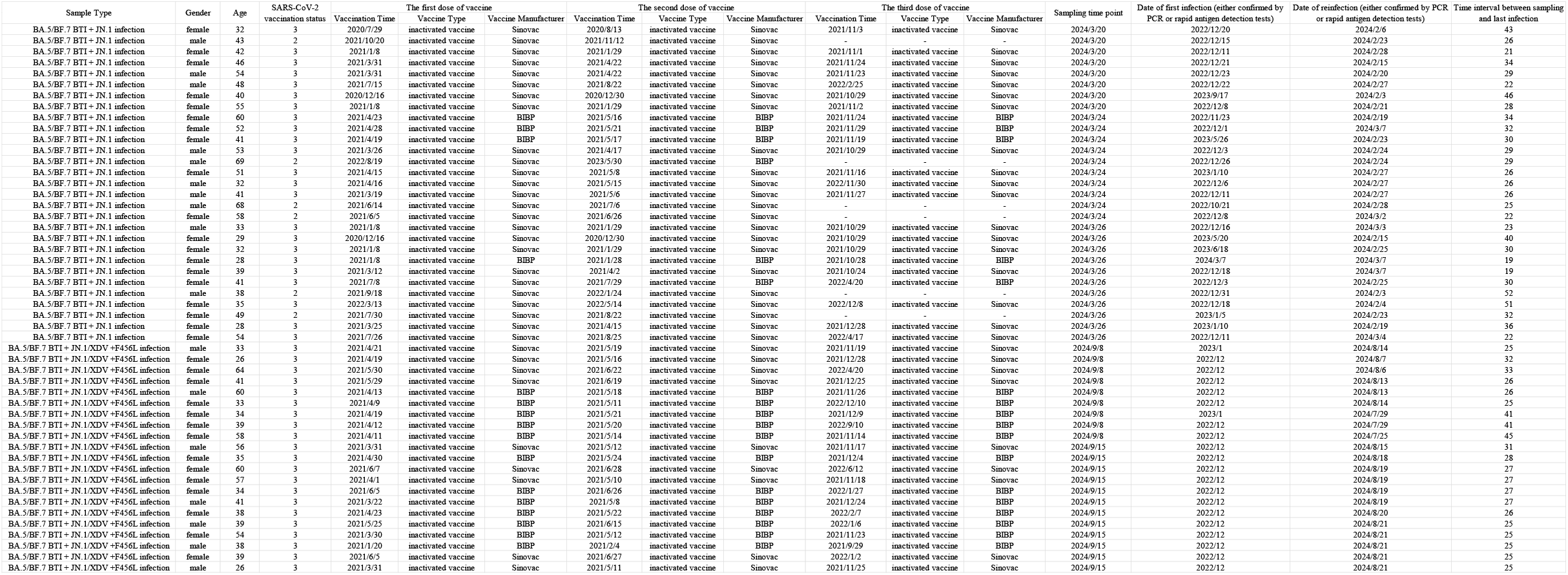
Information of SARS-CoV-2 convalescent patients participating in this study.

